# Bacteriophage cooperation suppresses CRISPR-Cas3 and Cas9 immunity

**DOI:** 10.1101/279141

**Authors:** Adair L. Borges, Jenny Y. Zhang, MaryClare F. Rollins, Beatriz A. Osuna, Blake Wiedenheft, Joseph Bondy-Denomy

## Abstract

>Bacteria utilize CRISPR-Cas adaptive immune systems for protection from bacteriophages (phages), and some phages produce anti-CRISPR (Acr) proteins that inhibit immune function. Despite thorough mechanistic and structural information for some Acr proteins, how they are deployed and utilized by a phage during infection is unknown. Here, we show that Acr production does not guarantee phage replication, but instead, infections fail when phage population numbers fall below a critical threshold. Failing infections can be rescued by related phages that act as Acr donors, demonstrating that infections succeed if a sufficient Acr dose is contributed to a single cell by multiple phage genomes. The production of Acr proteins by phage genomes that fail to replicate leave the cell immunosuppressed, which predisposes the cell for successful infection by other phages in the population. This “cooperative” phage mechanism for CRISPR-Cas inhibition demonstrates inter-virus cooperation that may also manifest in other host-parasite interactions.

## INTRODUCTION

Bacteria and the viruses that infect them (phages) are engaged in an ancient evolutionary arms race, which has resulted in the emergence of a diversity of CRISPR-Cas (clustered regularly interspaced short palindromic repeats and CRISPR-associated genes) adaptive immune systems (Koonin et al., 2017). CRISPR-Cas immunity is powered by the acquisition of small fragments of phage genomes into the bacterial CRISPR array, the subsequent transcription and processing of these arrays to generate small CRISPR RNAs, and the RNA-guided destruction of the phage genome (Barrangou et al., 2007; Brouns et al., 2008; Garneau et al., 2010; Levy et al., 2015). The destruction of foreign DNA by CRISPR-Cas has been shown to prevent the acquisition of plasmids, DNA from the environment, phage lytic replication, and prophage integration (Barrangou et al., 2007; Bikard et al., 2012; Cady et al., 2012; Edgar and Qimron, 2010; Garneau et al., 2010). In bacterial populations, these systems provide a fitness advantage to their host microbe during times of phage presence in the environment (van Houte et al., 2016; Westra et al., 2015).

To combat the potent action of RNA-guided CRISPR-Cas nucleases, phages have developed inhibitor proteins called anti-CRISPRs (Acrs). Acr proteins have been discovered in phages, prophages, mobile islands, and core genomes across many distinct bacteria and archaea (Borges et al., 2017; He et al., 2018; Pawluk et al., 2017). Specific Acr proteins inhibit the Type I-F, I-E, and I-D CRISPR-Cas3 have been identified (Bondy-Denomy et al., 2013; He et al., 2018; Pawluk et al., 2014; 2016b), and also other Acr proteins that inhibit Type II-A and II-C (Hynes et al., 2017; Pawluk et al., 2016a; Rauch et al., 2017) CRISPR-Cas9 systems. Phylogenetic studies indicate that these proteins are likely ubiquitous in coevolving populations of bacteria and phages (Pawluk et al., 2017) and provide a significant replicative advantage to phages in the presence of CRISPR immunity (van Houte et al., 2016).

Anti-CRISPRs were first identified in phages that neutralize the *Pseudomonas aeruginosa* type I-F system (<u>a</u>nti-<u>CR</u>ISPR type I-<u>F</u>, AcrIF1-5)(Bondy-Denomy et al., 2013), and five more I-F anti-CRISPRs (AcrIF6-10) were subsequently identified in various mobile genetic elements (Pawluk et al., 2016b). The I-F Csy surveillance complex (also called I-F Cascade) is comprised of four proteins (Csy1-4) and a *trans*-acting nuclease/helicase protein, Cas2/3 (Rollins et al., 2017; Wiedenheft et al., 2011). Anti-CRISPR proteins function by interacting directly with the Csy complex and inhibit DNA binding or bind to Cas2/3 and prevent nuclease-mediated degradation (Bondy-Denomy et al., 2015). The structures of type I-F Acr proteins AcrIF1, AcrIF2, and AcrIF3 have been solved in complex with their target proteins, revealing mechanistically distinct inhibitors that bind tightly to their target Cas protein (Chowdhury et al., 2017; Guo et al., 2017; Maxwell et al., 2016; Wang et al., 2016). Together with the recent identification and characterization of proteins that inhibit Cas9, a common theme in Acr function has been revealed; all characterized Acr proteins block phage DNA binding or cleavage (Dong et al., 2017; Harrington et al., 2017; Pawluk et al., 2016a; Rauch et al., 2017).

Despite a detailed mechanistic understanding, however, it is unknown how anti-CRISPR proteins protect a phage during infection. Where phage DNA cleavage has been assessed *in vivo*, it occurs in as little as 2 minutes (Garneau et al., 2010), suggesting that CRISPR attack may outpace the ability of *de novo* Acr synthesis and action. During phage infection, we hypothesized that successful inhibition of CRISPR-Cas immunity by Acr proteins would be challenging, as all components of the *P. aeruginosa* immune system are expressed prior to phage infection (Bondy-Denomy et al., 2013; Cady et al., 2012).

Here, we utilize the diverse array of AcrIF proteins encoded by phages to demonstrate that CRISPR-Cas inactivation is a challenge, and that the sufficient concentration of Acr proteins necessary to inactivate CRISPR-Cas is contributed by multiple phage genomes. While initial phage infections may fail, due to the rapid action of the CRISPR-Cas system, immunosuppression can result due to the production of Acr proteins, which leads to a population-level benefit for other phages. We propose that pathogens can contribute to the “remodeling” of their host cell via rapid protein production, even if the initial infecting genomes are cleared, opening the door for their clones.

## RESULTS

### Anti-CRISPR proteins are imperfect CRISPR-Cas inhibitors

We utilized the diversity of *acr* genes encoded by phages infecting *P. aeruginosa* to determine the mechanism of CRISPR-Cas neutralization during infection. Five natural phages, each encoding a single *acrIF* gene, were selected to represent *acrIF1-IF4* and *acrIF7* (*acrIF5* does not exist as the sole *acrIF* gene on any phage, *acrIF6, F8-F10* are not encoded by this phage family). Three of the five phages exhibited reduced efficiency of plaquing (EOP) on *P. aeruginosa* strain PA14, possessing a type I-F CRISPR-Cas system (Figure 1A, WT:pEmpty normalized to plaquing on ΔCRISPR). Overexpression of a targeting crRNA (WT:pSp1) exacerbated anti-CRISPR inefficiency, limiting the replication of all anti-CRISPR phages by at least one order of magnitude. This suggests that anti-CRISPR proteins are unable to fully protect their associated phage genome.

**Figure 1.**
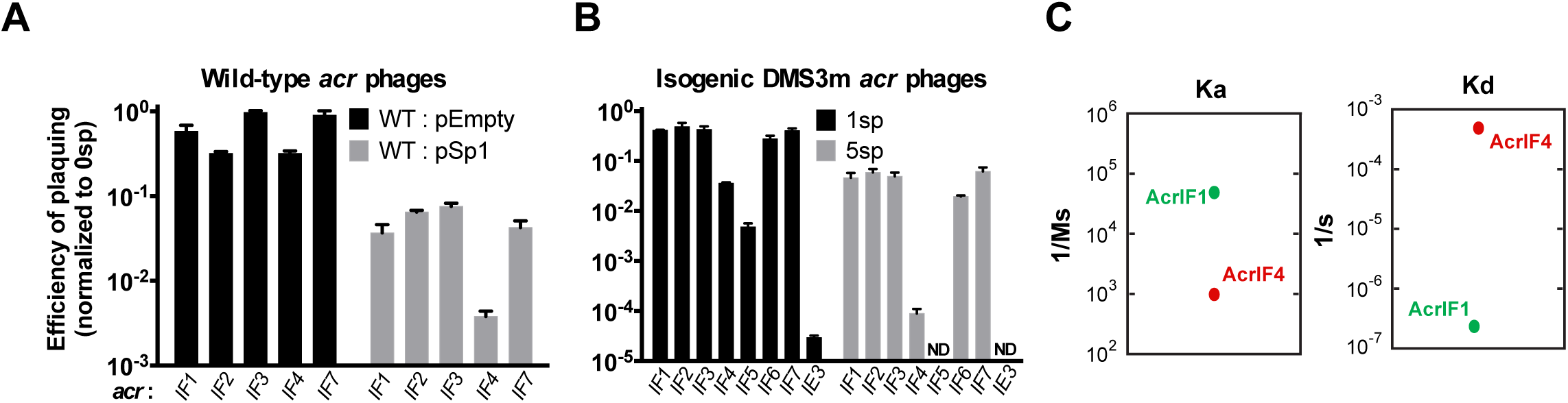
Anti-CRISPRs are imperfect CRISPR-Cas inhibitors. **(A)** Efficiency of plaquing (EOP) of 5 related phages bearing distinct *acrIF* genes (JBD30_*acrIF1*_, MP29_*acrIF2*_, JBD88a_*acrIF3*_, JBD24_*acrIF4*_, LPB1_*acrIF7*_) on *Pseudomonas aeruginosa* strain PA14. Plaque forming units (PFUs) were quantified on wild-type PA14 with 1-2 natural targeting spacers (WT + pEmpty) or on PA14 overexpressing 1 targeting spacer (WT + pSp1), then normalized to the number of PFUs measured on a non-targeting PA14 derivative (0sp). Data are represented as the mean of 3 biological replicates +/-SD. **(B)** EOP of isogenic DMS3m phages with *acrIF1-7* or *acrIE3* in the DMS3m *acr* locus. EOP was calculated as PFU counts measured on WT PA14 with 1 targeting spacer (1sp) or a laboratory evolved PA14 derivative with 5 targeting spacers (5sp) normalized to PFU counts measured on non-targeting PA14 (0sp). Data are represented as the mean of 3 biological replicates +/-SD. ND, not detectable. **(C)** Plot of association (Ka) and dissociation (Kd) rates for AcrIF1 (data adapted from Chowdhury et al. 2017) and AcrIF4 binding the PA14 Csy complex. See Figure S2 for SPR AcrIF4 sensogram.

To assess anti-CRISPR strength directly, an isogenic phage panel was generated by replacing the *acrIE3* gene of phage DMS3m with single *acrIF* genes *F1-F7* (DMS3m_acrIF1_-DMS3m_acrIF7_, Figure S1). WT PA14 (1 spacer targeting DMS3m, “1sp”) and a laboratory evolved strain (5 spacers targeting DMS3m, “5sp”) were challenged with the recombinant phages, revealing that all anti-CRISPRs are imperfect during infection (Figure 1B). For phages encoding *acrIF1, F2, F3, F6* or *F7*, >90% of phage in the population failed to replicate (EOP=10^-1^) when faced with 5 targeting spacers. *acrIF4* and *F5* were very weak, with 99.0-99.99% of phages failing to replicate, depending on the CRISPR spacer content. We conclude that phages encoding *acrs* remain sensitive to CRISPR-Cas immunity, suggesting that anti-CRISPR deployment and action is an imperfect process.

The observation above identified groups of “strong” and “weak” Acr proteins. We selected one representative from each group, and a third Acr that does not target the I-F CRISPR system (i.e. AcrIE3), as a negative control. AcrIF1 was selected as a model strong inhibitor, as its mechanism and binding affinity are known (Csy complex binding, K_D_ = 2.5 x 10^-11^ M (Bondy-Denomy et al., 2015; Chowdhury et al., 2017)). In contrast, AcrIF4 is a weak inhibitor that also binds the Csy complex (Bondy-Denomy et al., 2015), but with a significantly slower on-rate and faster off-rate compared to AcrIF1 (Figure 1C, Figure S2).

We next assessed the survival of bacterial cell populations over time, when infected with phages that must deploy Acr proteins, but where apparently only a minority of the population actually replicates. Phages were locked in the lytic cycle for the purpose of this experiment, by knocking out the *C repressor* gene (*gp1*) in DMS3m_*acrIF1*_, DMS3m_*acrIF4*_, and DMS3m_*acrIE3*_, to enable the tracking of bacterial survival over time. The virulent (vir) phages were used to infect 5sp strains in liquid culture, and bacterial growth measured. Given the >4 order of magnitude difference in the KD of AcrIF1 and AcrIF4 for their binding partner, we reasoned that different phage concentrations may be required to inactivate CRISPR-Cas function. In the presence of CRISPR immunity, bacterial death only occurred at multiplicities of infection (MOI, input plaque forming units per colony forming unit) greater than 0.02 for the strong *acrIF1* (Figure 2A) and greater than an MOI of 2.0 (≥10^7^ PFU) for the weak *acrIF4* (Figure 2B). Despite this high MOI-dependence, no selection or evolution of phage DMS3m_*acrIF4*_ occurred throughout the experiment, as output phages remained as sensitive to CRISPR-Cas immunity as the input population (Figure S3). The phage encoding *acrIE3* had no impact on bacterial survival when faced with CRISPR immunity (Figure 2C), while in the absence of CRISPR, phages at all concentrations cleared bacterial cultures (Figure 2D-2F). These data demonstrate for bacterial lysis to occur, a critical phage concentration threshold exists that is inversely proportional to Acr strength.

**Figure 2.**
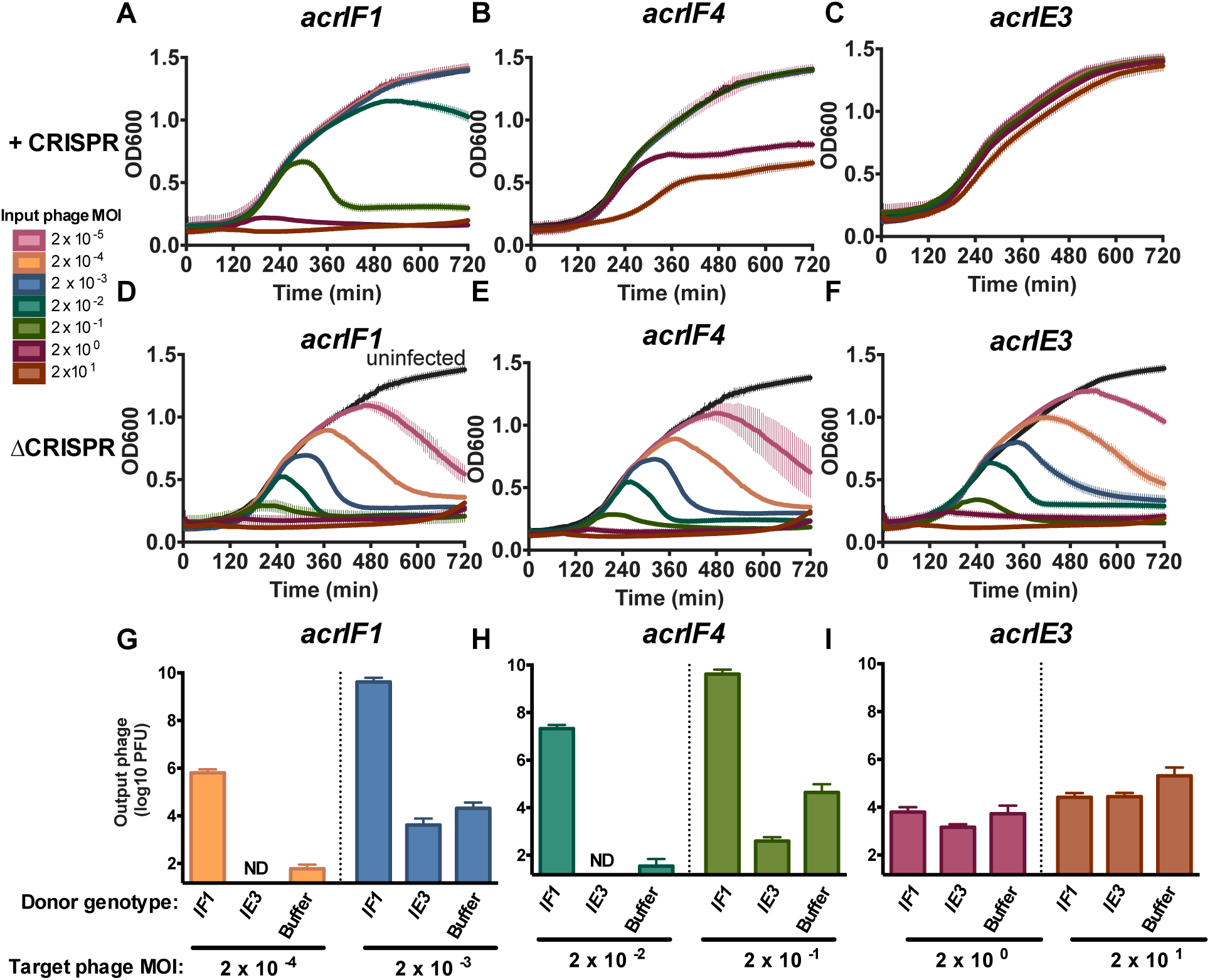
Anti-CRISPR success requires cooperative infections during lytic growth. **(A-F)** 12 h growth curves of *P. aeruginosa* strain PA14 with 5 targeting spacers (+CRISPR, panels **A-C**) or no CRISPR-Cas function (ΔCRISPR, **D-F**) infected with virulent variants of DMS3m_*acrIF1*_, DMS3m_*acrIF4*_, or DMS3m_*acrIE3*_ at multiplicity of infection (MOI) increasing in 10-fold steps from 2x10^-5^ to 2x10^1^ (rainbow colors) or uninfected (black). Colors correspond to the MOI legend and growth curves. OD600 is represented as the mean of 3 biological replicates +/-SD. ND, not detectable. **(G-I)** Replication of virulent DMS3m_*acr*_ phages (target phage) in the presence of 10^6^ PFU (MOI 0.2) hybrid phage (donor) in PA14 with 5 targeting spacers expressing the JBD30 C-repressor. Phages were harvested after 24 hours of co-culture and DMS3m_*acr*_ phage PFUs were quantified on PA14 0sp expressing the JBD30 C repressor. Phage output is represented as the mean of 3 biological replicates +/-SD. ND, not detectable.

### Lytic replication requires a critical Acr protein concentration

Concentration-dependent success of CRISPR-Cas neutralization during phage infection could be caused by an excess of phage DNA targets that overwhelm a cell by titrating away the Csy complexes, or by the contribution of Acr proteins from multiple phage genomes in a single cell. To experimentally address these models, we genetically modified phages to render them non-replicative, thus allowing the independent titration of a subset of phages in the population. The C repressor gene (gp1) and surrounding immunity region from phage JBD30 was introduced into DMS3m phages, generating a hybrid phage (Figure S4A), whose replication could be prevented by the overexpression of the JBD30 C repressor (*gp1*, Figure S4B). The JBD30 C repressor does not impact the replication of DMS3 phages, however, as they are from different immune families (heteroimmunity). In the presence of the JBD30 C repressor, high doses of the hybrid phage did not replicate, confirming the efficacy of *gp1* overexpression (Figure S4C). This enabled the mixing of two independent phage populations, where the replication of one could be experimentally blocked, rendering it a sacrificial genome and Acr “donor”. This was used to assess the Acr concentration dependency model.

In the presence of sacrificial donor phages encoding AcrIF1, we observed a striking contribution to CRISPR-Cas neutralization. This enabled DMS3m_acrIF1_ (Figure 2G) and DMS3m_acrIF4_ (Figure 2H) to replicate from input MOIs that are unsuccessful in the absence of the donor phage (Figure 2H,2G see “buffer”). Although the input concentration of donor phages was modest (10^6^ PFU, MOI = 0.2), this enabled the replicative phage to achieve an increase of titers by 4-5 orders of magnitude. However, this helping effect was not observed when the replicative phage did not encode an AcrIF protein (Figure 2I), or when the donor phage produced *acrIE3* (Figure 2H-2I). Additionally, any potential lysogens formed by the DMS3m donor phage in this experiment would not have amplified the replicating phage, as these lysogens remain resistant to superinfection (Figure S4D). These data demonstrate that Csy complex titration by infecting phage genomes alone does not appear to be a significant “anti-CRISPR” factor in the outcome of phage replication or bacterial survival. Instead, the determinant of phage replicative success is the concentration of Acr proteins reached in single cells, which is achievable by Acr production from independent phage genomes. This suggests that multiple phage genomes can contribute to the “immunosuppression” of a single host by contributing separate doses of Acr protein, potentially explaining the observed concentration thresholds and inefficiencies observed in plaquing measurements.

### Lysogeny requires Acr proteins contributed by transient intracellular genomes

All phages encoding Acr proteins that infect *P. aeruginosa* are naturally temperate, and can form lysogens by integrating into the bacterial genome. We therefore measured the impact of CRISPR and Acr proteins on lysogeny establishment during a single round of infection. While previous experiments examined cumulative phage replication in the lytic cycle over many hours, assaying lysogen formation over a short time frame is ideal for understanding the initial events that determine phage genome survival or cleavage. Additionally, lysogeny provides a direct readout for phage genome survival (i.e. an integrated prophage), while in lytic replication, phage survival leads to a dead cell that cannot be recovered. For these experiments, we selected the weak AcrIF4 protein as it provided the largest dynamic range of inefficiency in a single round of infection.

We generated derivatives of DMS3m_*acrIF4*_ and DMS3m_*acrIE3*_ marked with a gentamicin resistance cassette at the end of the genome, replacing a nonessential gene, *gp52*. This allowed the independent titration of two distinct replication-competent phage populations and the selection and analysis of stable lysogens after the experiment. These phages were used to infect ΔCRISPR cells (0sp) for a time span less than a single round of infection (50 minutes, Figure S5), and the number of gentamicin resistant lysogens was assessed. In the absence of CRISPR selection, a linear increase in the number of lysogens with increasing MOI was observed, over ∼4 orders of magnitude (Figure 3A-3B, circles). In the presence of spacers targeting DMS3m (5sp), CRISPR reduced the efficiency of lysogeny (EOL, compared to 0sp) for the weak *acr* phage DMS3m_acrIF4_, to such a degree that lysogeny was undetectable at low MOIs, and impaired by 100-1000-fold at higher MOIs (Figure 3A, triangles). DMS3m_acrIF4_ demonstrated concentration dependence for successful lysogeny, with EOL values below or at the limit of detection for lower MOIs, increasing to EOL = 0.01 at higher MOIs (Figure 3C). Phage DMS3m_acrIE3_ formed no lysogens at all input concentrations tested, demonstrating that Acr-mediated immune suppression is required to establish lysogeny (Figure 3B, 3D).

**Figure 3.**
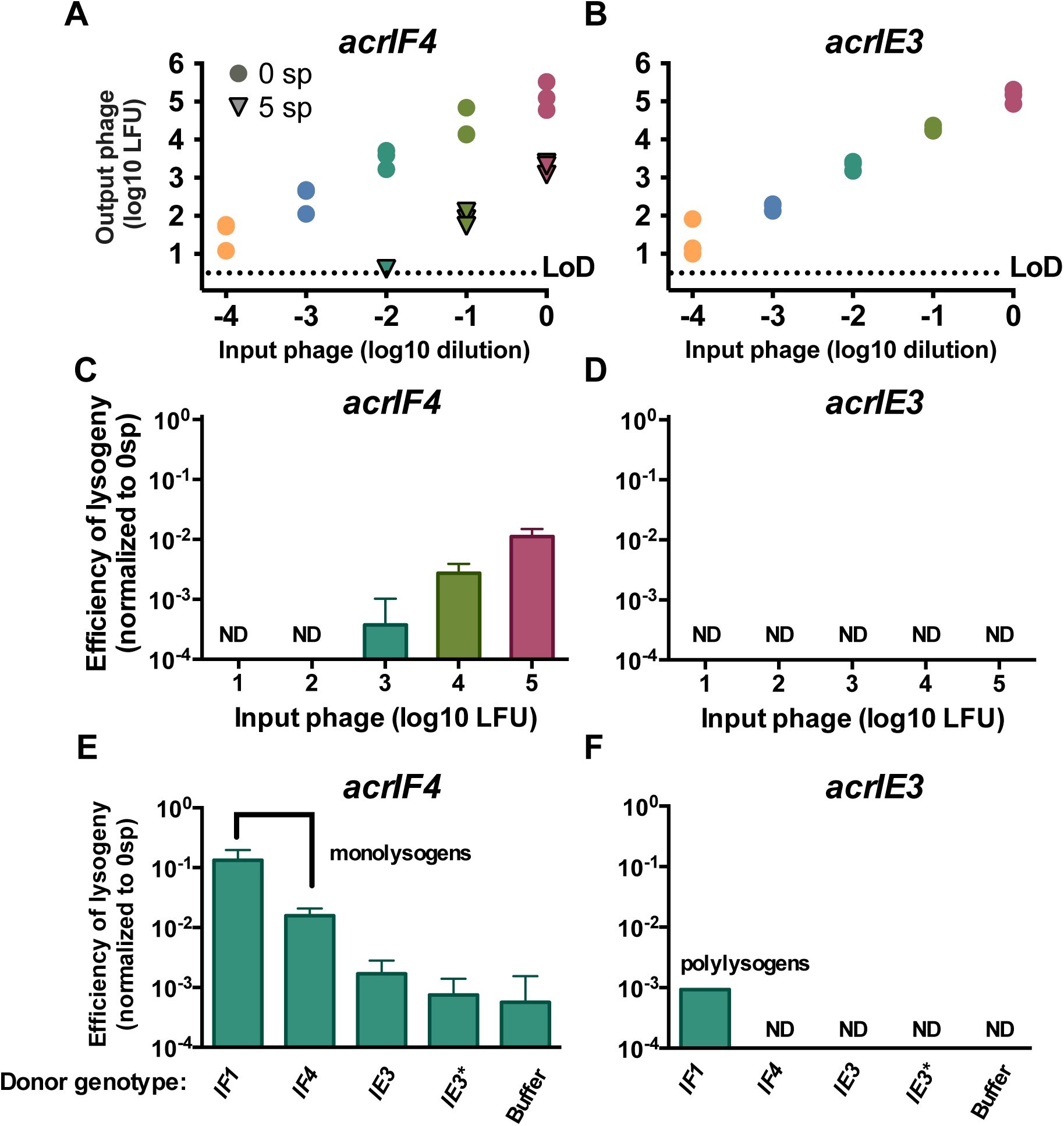
Immunosuppression facilitates acquisition of a marked prophage. (**A,B**) Acquisition of marked DMS3m_*acrIF4*_ or DMS3m_acr*IE3*_ prophage by PA14 with 0 spacers (0 sp, circles) or 5 targeting spacers (5 sp, triangles). This experiment was performed in biological triplicate, and individual replicate values are displayed. LoD, limit of detection. **(C,D)** Efficiency of lysogeny (EOL) of DMS3m_*acrIF4* *gp52∷gentR*_ and DMS3m_*acrIE3* *gp52∷gentR*_ in the presence of CRISPR targeting. EOL was calculated by dividing the output lysogens forming units (LFUs) from the strain with 5 targeting spacers (5sp) to the number of LFUs in PA14 with 0 targeting spacers (0sp). Data are represented as the mean of 3 biological replicates +/-SD. ND, not detectable. **(E,F)** EOL of 10^3^ LFUs of DMS3m_*acrIF4* *gp52∷gentR*_ and DMS3m_*acrIE3* *gp52∷gentR*_ in the presence of 10^7^ PFU of the indicated DMS3m_*acr*_ phage. Data are represented as the mean of 3 biological replicates +/-SD. ND, not detectable. See Figure S6 for lysogen genetic analysis.

We hypothesized that Acr concentration dependence for CRISPR neutralization during lysogeny could also be explained by phage cooperation, and that below threshold concentrations of DMS3m_*acrIF4*_ could be rescued by the addition of Acr donor phages *in trans*. To test this hypothesis, we infected the 5sp strain with a mixture of 10^3^ LFU DMS3m_*acrIF4*_ *gp52∷gent* and 10^7^ PFU of unmarked Acr donor phages, and measured the EOL of DMS3m_*acrIF4*_ *gp52∷gent*. The EOL of DMS3m_*acrIF4*_ *gp52∷gent* increased by 2 orders of magnitude with Acr donor phage DMS3m_*acrIF1*_, while DMS3m_acrIF4_ donor phage increased EOL by 1 order of magnitude (Figure 3E). No such increase in lysogeny was observed for a phage without an *acrIF* gene, DMS3m_*acrIE3*_ *gp52∷gent* (Figure 3F). The addition of Acr donor phages DMS3m_*acrIE3*_, or an escaper phage DMS3m_acrIE3_* had no effect on the EOL of the marked DMS3m_*acrIF4*_ phage, demonstrating that the donor phage must be an Acr-producer.

To determine the specific mechanism of anti-CRISPR donation leading to survival DMS3m_*acrIF4*_ *gp52∷gent*, we used the resulting lysogens as a genetic record of infection success for both the marked phage and the unmarked phage. This family of Mu-like phages integrates randomly into the host genome, and therefore strains with multiple prophages are possible (Bondy-Denomy et al., 2016). We assayed the lysogens resulting from the experiment described above for the presence of the donor phage genome in addition to the DMS3m_*acrIF4*_ *gp52∷gent* prophage. All resulting lysogens tested (n=48) possessed only the marked prophage, with none possessing the Acr donor prophage (Figure S6). This demonstrates that the transient presence of the Acr donor phage genome (i.e. no lysogeny) in the same cell was sufficient to generate enough Acr protein to protect the marked phage, leading to the establishment of lysogens that would not exist if not for the Acr donor (Figure 3E, compare “buffer” to “IF1”). The only time rare double lysogens were isolated, was when the marked phage DMS3m_*acrIE3*_ *gp52∷gent* was used with the unmarked DMS3m_*acrIF1*_ Acr donor phage (Figure 3F), since the prophage encoding *acrIE3* would be unable to neutralize CRISPR-Cas after lysogenic establishment and long-lasting protection from the donor prophage was required. Collectively, these data are consistent with a model where the production of Acr proteins from a transient phage genome prior to its cleavage generates an immunosuppressed cell that can be successfully parasitized by another phage upon re-or co-infection(s).

### Stoichiometric inhibitors of Cas9 required bacteriophage cooperation

The intrinsic inefficiency of stoichiometric inhibitors is likely due to the requirement for the rapid synthesis of a high concentration of inhibitors before phage genome cleavage. To determine whether this model generally applies to other stoichiometric inhibitors of bacterial immunity, we engineered a *P. aeruginosa* strain to express the Cas9 protein from *Streptococcus pyogenes* (SpyCas9) and a DMS3m phage to express a previously identified Cas9 inhibitor, AcrIIA4 (Dong et al., 2017; Rauch et al., 2017). With this entirely heterologous system, we again observed inefficiency for a phage relying on an Acr protein. Spot-titration of phage lysates on a strain expressing a single guide RNA (sgRNA) targeting DMS3m decreased the titer of DMS3m_acrIE3_ by many orders of magnitude, while DMS3m_acrIIA4_ was protected (Figure 4A). However, quantification of the efficiency of plaquing again revealed that relying on an Acr protein for replication is imperfect, with an EOP = 0.4 (Figure 4B). In lytic replication infection experiments, DMS3m_acrIIA4_ displayed concentration-dependent bacterial lysis in the presence of CRISPR targeting (Figure 4C), while infection with DMS3m_acrIE3_ did not affect bacterial growth (Figure 4D). In the absence of CRISPR-Cas targeting, however, both phages killed their hosts at all phage concentrations tested (Figure 4E, 4F). To determine whether this concentration dependence for Cas9 inhibition was also a result of insufficient intracellular Acr dose, a non-replicative hybrid DMS3m_AcrIIA4_ phage was generated and used as a donor during infection. Indeed, increased delivery of AcrIIA4 to cells allowed DMS3m_acrIIA4_ to replicated robustly from a concentration that was previously unsuccessful (Figure 4G), demonstrating phage cooperation to neutralize CRISPR-Cas9. Unlike CRISPR-Cas3 experiments described previously, where 5 spacers were targeted to the DMS3m phage, here a single spacer sequence was used, which allowed for phage escape mutations to occur. Because of this, we observed that DMS3m_acrIE3_ could also be helped by increased doses of AcrIIA4 from the phage population (Figure 4H), likely protecting the genome *in trans*, enabling escaper evolution.

**Figure 4.**
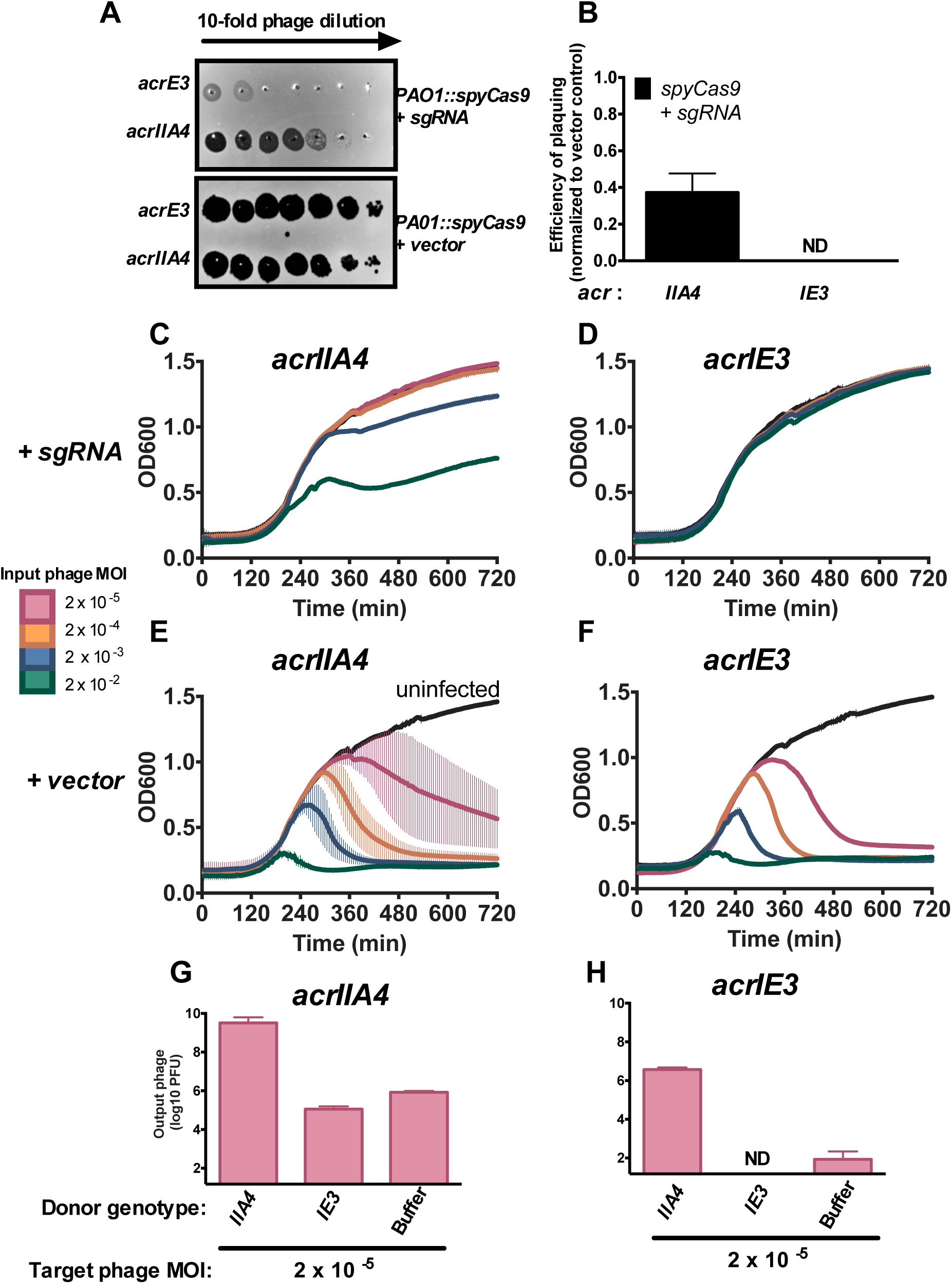
Cas9 anti-CRISPR AcrIIA4 requires cooperative infection to neutralize Type II-A CRISPR immunity. **(A)** 10-fold serial dilutions of DMS3m_*acrIE3*_ or DMS3m_*acrIIA4*_ plated on a lawn of *Pseudomonas aeruginosa* strain PAO1 expressing *Streptococcus pyogenes* Type II-A Cas9 (PAO1∷SpyCas9) and single guide RNA (+ sgRNA) or non-targeting control (+ vector). **(B)** Efficiency of plaquing of DMS3m_*acrIIA4*_ and DMS3m_*acrIE3*_ was calculated by normalizing PFU counts on a targeting strain of PAO1∷SpyCas9 (+sgRNA) to PFU counts on a non-targeting strain of PAO1∷SpyCas9 (+vector). Data are represented as the mean of 3 biological replicates +/-SD. ND, not detectable. Δ **(C-F)** 12 hour growth curves of PAO1∷spyCas9 expressing a targeting sgRNA (+ sgRNA, panels **C-D**) or a non-targeting vector control (+vector, E-F) that were infected with virulent DMS3m_*acrIIA4*_ or DMS3m_*acrIE3*_ at multiplicities of infection (MOI, rainbow colors) from 2x10^-5^ to 2x10^-2^. Growth curves of uninfected cells are shown in black. OD600 values are represented as the mean of 3 biological replicates +/-SD. **(G-H)** Replication of virulent DMS3m_*acr*_ phages (target phage) in the presence of 10^7^ PFU (MOI 2) hybrid phage (donor) in PAO1∷SpyCas9 + sgRNA expressing the JBD30 C-repressor. Phages were harvested after 24 hours and DMS3m_*acr*_ phage PFUs quantified on PAO1∷SpyCas9 + vector expressing the JBD30 C repressor. Phage output is represented as the mean of 3 biological replicates +/-SD. ND, not detectable.

## DISCUSSION

Here we demonstrate that the necessary intracellular concentration of an anti-immunity protein to achieve inactivation of cellular immunity depends on the relative strengths of both the inhibitor and the immunity pathway, which dictates the number of infecting viruses required in the population. We conclude that a single cell can become immunosuppressed by anti-immune protein contributions from independent infection events. In the absence of viral replication, these infection events serve to contribute to the inactivation of cellular immunity, thus enhancing the probability of successful infection events in the future (Figure 5). We expect that cooperation of this sort is necessary when the immune process acts rapidly and irreversibly on the infecting viral genome, as CRISPR-Cas immunity does.

**Figure 5.**
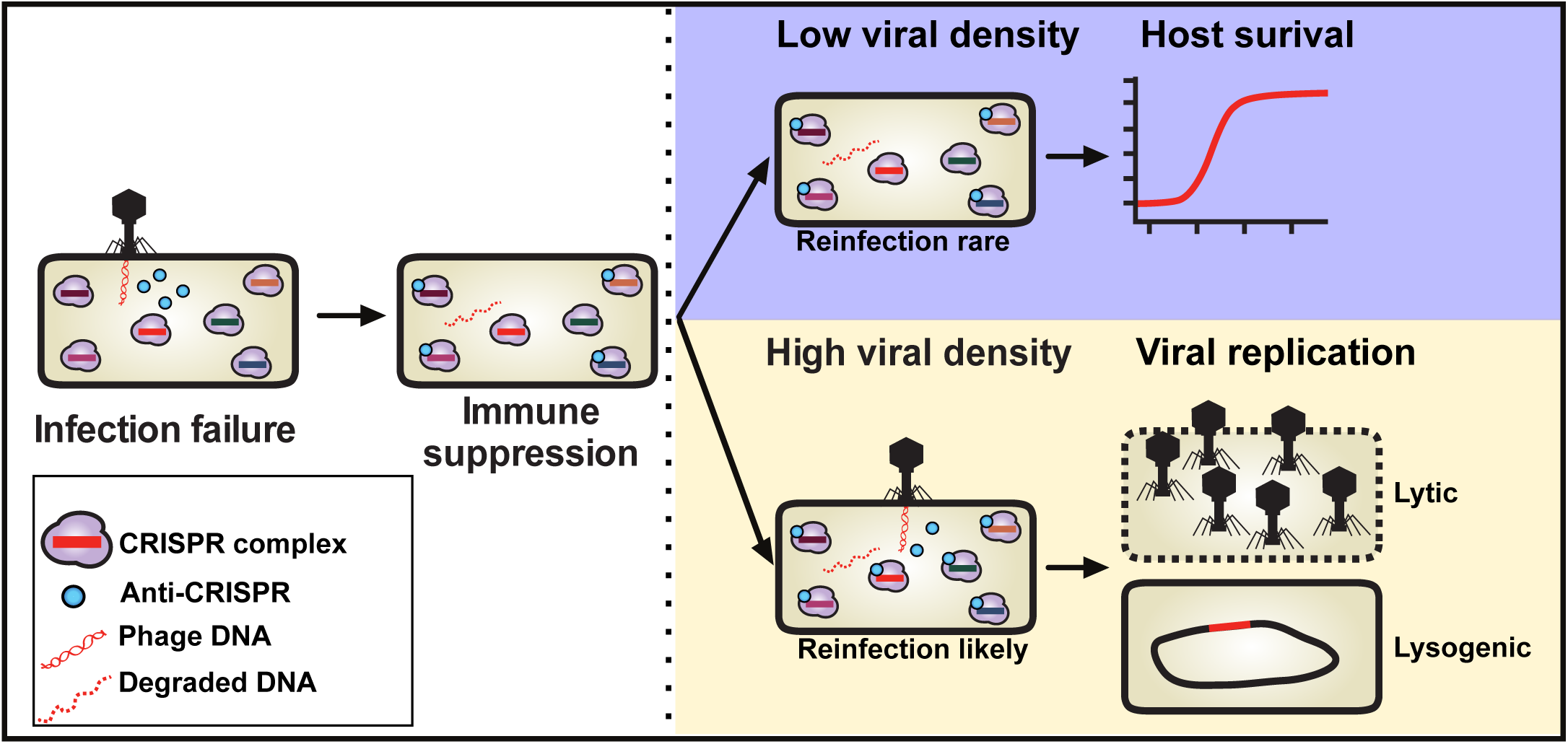
Phage cooperation to suppress CRISPR-Cas immunity. *Left*: Failed infections generate an immunosuppressed cell. While the phage genome may be cleaved (dashed red line), Acr protein (small cyan circles) will be produced, binding to some CRISPR-Cas complexes (larger purple structures, with crRNA). *Right:* At low parasite density, it is unlikely the cell will be re-infected and the host will survive. As high parasite density, the likelihood that the immunosuppressed cell with be re-infected is higher. Co-or re-infection will lead to successful parasite replication and amplification of the infectious population.

The ability of phages to replicate in the lytic cycle or establish lysogens is impacted dramatically by the number of phages in the population. To demonstrate phage-phage cooperation for the deployment of Acr proteins, three distinct genetic strategies were used, allowing the independent titration of phages encoding a defined Acr protein: i) non-replicative Acr donor phages, ii) marked and unmarked phages to follow the fate of only one phage, and iii) the prophage status of lysogens, as a genetic record of phage success. By restricting the replication of a subset of phages, we could track the lytic replication of a low-dose of wild-type phages in the presence of DNA target or Acr donors to determine the molecular determinants of density-dependent phage replication. In contrast, the prophage acquisition experiment allows a more natural scenario, where all phages are capable of replication, but an antibiotic resistance marker allows us to track the outcome of one phage genotype, starting from a concentration below its threshold for successful lysogeny. This demonstrated that wild-type Acr-donor phages in the population contribute to cellular immunosuppression, enabling the formation of lysogens that did not occur in their absence. The presence of only a single, marked prophage in the bacterial genome proves that the donor phage neither entered the lytic cycle (this would kill the cell), nor lysogenized (prophage would be present), but had been present in the cell transiently.

The key result here is the observation that phages can remodel their host cell, even in the absence of a replicating or integrated genome. It has long been known that integrated prophages modulate host phenotypes via gene expression, including superinfection exclusion, toxin production, and the production of Acr proteins (Bondy-Denomy et al., 2013; Bondy-Denomy and Davidson, 2014; Bondy-Denomy et al., 2016; Waldor and Mekalanos, 1996; Weigle and Delbruck, 1951). Although less commonly described, there is also precedent for lytic phages to impact an interaction with other phages during replication. The Imm protein produced by the phage T4 prevents other phages in the environment from infecting the cell that one phage is currently replicating within (Lu and Henning, 1989). This has been attributed to preventing superinfection and the potential disruption of the carefully timed phage replication cycle. This supports a conclusion that co-and/or re-infections are a common occurrence in nature. However, here, we propose a new model of phage-induced host remodeling, whereby a transient-unsuccessful infection produces proteins that inactivate defense, enabling future infections. This altruism would be evolutionarily advantageous to phage clones within the population, where all are providing Acr proteins to neutralize CRISPR.

A distinct, but notable observation from this work is that not all Acr proteins operate at equivalent strengths. However, encoding even a weak inhibitor (e.g. AcrIF4). still provides a significant advantage to the phage, compared to lacking them entirely. We show that AcrIF4 binds the Csy complex with affinities that are orders of magnitude weaker than Acr proteins like AcrIF1. We selected AcrIF1 as a model strong Acr protein because of its comparable mechanism of action to AcrIF4 (i.e. Csy complex binding), and consider it representative of other strong Acr proteins (AcrIF2, F3, F6, F7), based on EOP data. Going forward, we speculate that the strongest Acr proteins would be enzymatic in nature, allowing rapid and efficient inactivation of CRISPR complexes in a sub-stoichiometric manner, although no such Acr mechanism has been discovered. While not an enzyme, the recent demonstration of the AcrIIC3 protein inactivating two Cas9 proteins at the same time would likely be a more efficient path towards CRISPR neutralization (Harrington et al., 2017).

The challenge of neutralizing a pre-expressed CRISPR-Cas system likely explains why stoichiometric inhibitors like Acr proteins are imperfect, and phages relying on them are partially targeted by CRISPR. The sacrificial, population-level aspect of CRISPR inhibition is reminiscent of the manifestations of CRISPR adaptation in populations of bacterial cells. The majority of infected naïve host cells die, before a clone with a new spacer emerges (Barrangou et al., 2007; Hynes et al., 2014). In the case of anti-immunity, many phages die in order to inhibit CRISPR on a single cell level, and this must happen at sufficient frequency within an ecosystem for phage to prevail. We suspect that this mechanism of cellular immunosuppression and inter-parasite cooperation may have parallels in other host-pathogen interactions, where concentration dependence manifests at predictable levels due the strengths of immune and anti-immune processes.

## SUPPLEMENTAL INFORMATION

Figures S1-S6 are provided separately.

## AUTHOR CONTRIBUTIONS

A.L.B. and J.B.D. designed experiments and wrote the manuscript. A.L.B. engineered phage variants and conducted all experiments with the exception of binding affinity measurements. J.Y.Z. assisted with experimental design and execution, phage engineering, and data collection. B.A.O. built the Cas9 strain in *Pseudomonas aeruginosa.* J.B.D. supervised experiments. M.R. conducted Acr binding affinity measurements under the supervision of B.W. All authors contributed to data analysis and edited the manuscript.

## ACKNOWLEDGEMENTS

The Bondy-Denomy lab was supported by the University of California San Francisco Program for Breakthrough in Biomedical Research, funded in part by the Sandler Foundation, and an NIH Office of the Director Early Independence Award (DP5-OD021344). Research in the Wiedenheft lab is supported by the National Institutes of Health (P20GM103500, P30GM110732, R01GM110270, R01GM108888 and R21 AI130670), the National Science Foundation EPSCoR (EPS-110134), the M. J. Murdock Charitable Trust, a young investigator award from Amgen, and the Montana State University Agricultural Experimental Station (USDA NIFA). The laboratory evolved PA14 strain with 5 spacers targeting DMS3 was generously provided by Stineke van Houte and Edze Westra (University of Exeter).

## METHODS

### Strains and growth conditions

*Pseudomonas aeruginosa* strains (UCBPP-PA14, PAO1) and *Escherichia coli* strains were cultured on lysogeny broth (LB) agar or liquid media at 37C. LB was supplemented with gentamicin (50 µg/mL for *P. aeruginos*a, 30 µg/mL for *E. coli*) to maintain the pHERD30T plasmid or carbenicillin (250 µg/mL for *P. aeruginos*a, 100 µg/mL for *E. coli*) to maintain pHERD20T or pMMBHE. To maintain pHERD30T and pMMBHE in the same strain of *P. aeruginosa*, double selection of 30 µg/mL gentamicin and 100 µg/mL carbenicillin was employed. In all *P. aeruginosa* experiments, expression from pHERD20/30T was induced with 0.1%

### PAO1 SpyCas9 expression strain

SpyCas9 expressed from the P_LAC_ promoter of pUC18T-mini-Tn7T-Gm was integrated into the *P. aeruginosa* strain PAO1 chromosome by electroporation and Flp-mediated marker excision as previously described (Choi and Schweizer, 2006). To generate the heterologous Type II-A PAO1 strain the PAO1-attTn7∷pUC18T-miniTn7T-P_LAC_-SpyCas9 strain was transformed with pBAO72 (pMMB67HE-P_LAC_-sgRNA) by electroporation. In all experiments with this strain, SpyCas9 and the sgRNA were induced with 1mM IPTG.

### Construction of recombinant DMS3m_*acr*_ phages

DMS3m_*acrIF1*_ was generated as in (Bondy-Denomy et al., 2013) by infecting cells containing a recombination plasmid bearing JBD30 genes 34-38 (the anti-CRISPR locus with large flanking regions). JBD30 naturally carries *acrIF1* and has high genetic similarity to DMS3m_*acrIE3*_, permitting for the selection of recombinant DMS3m phage which had acquired *acrIF1*. To generate the extended panel of DMS3m_*acr*_ phages in this work, recombination cassettes were generated with regions from up and downstream the anti-CRISPR genes from JBD30 Gibson assembled to flank the *acr* gene of interest on pHERD20T or pHERD30T (see S1 for *acr* gene sources and exact details of *acr* locus architecture). Recombinant phages were generated by infecting cells bearing these recombination substrates. DMS3m_*acr*_ phages were screened for their ability to resist CRISPR targeting, and the insertion of the anti-CRISPR gene was confirmed by PCR. Virulent derivatives of DMS3m_*acr*_ phages were constructed by deleting gp1 (C repressor) using materials and methods from Cady et al., 2012.

### Construction of DMS3m_*acr gp52∷GentR*_ phages

A recombination substrate with a gentamicin resistance cassette flanked by homology arms matching the DMS3m genome up and downstream of gp52 (450 bp and 260 bp, respectively) was assembled into pHERD20T using Gibson assembly. This recombination cassette was transformed into PA14 ΔCRISPR lysogenized with either DMS3m_*acrIE3*_ or DMS3m_*acrIF1*_. These transformed lysogens were grown under gentamicin selection for 16 h, then sub-cultured 1:100 into LB with gentamicin and 0.2 µg/mL mitomycin C to induce the DMS3m_*acr*_ prophage. Supernatants were harvested after 24 hours of induction, and used to infect PA14 ΔCRISPR for 24 h. These cells were then plated on gentamicin plates to select for cells that had acquired a prophage bearing the gentamicin resistance cassette, and gentamicin resistant lysogens were then re-induced with 0.2 µg/mL mitomycin C to recover the recombinant phage.

### Construction of DMS3m_*acr gp1-JBD30*_ (Hybrid_*acr*_) phages

DMS3m_*acrIE3*_ and JBD30_*acrIE3*_ were used to co-infect PA14 ΔCRISPR and the infected cells were mixed with molten top agar and poured onto solid plates. After 24 hours of growth at 30C, the phages were harvested by flooding the plate with SM buffer and collecting and clarifying the supernatant. Phages were then used to infect PA14 ΔCRISPR expressing the DMS3 C repressor from pHERD30T, and the infections were mixed with molten top agar and poured onto solid plates. After 24 hours of growth at 30°C, individual plaques with DMS3 morphology were picked, purified 3x by passage in PA14 ΔCRISPR and screened as shown in Figure S4B.

### Phage (PFU) quantification

Phage plaque forming units (PFU) were quantified by mixed 10µl of phage at a with 150 µl of an overnight culture of host bacteria. The infection mixture was incubate at 37 °C for 10 minutes to promote phage adsorption, then mixed with 3 mLs molten top agar and spread on an LB agar plate supplemented with 10 mM MgSO_4_. After 16 hours of growth 30 °C, PFUs were quantified.

### Phage titering

A bacterial lawn was generated by spreading 3 mLs of top agar seeded with 150 µl of host bacteria on a LB agar plate supplemented with 10 mM MgSO_4_. 3 µl of phage serially diluted in SM buffer was then spotted onto the lawn, and incubated at 30 °C for 16 hours.

### Measurement of anti-CRISPR binding kinetics by surface plasmon resonance

Purified Csy complex was covalently immobilized by amine coupling to the surface of a carboxymethyldextran-modified (CM5) sensor chip. Purified 6his-tagged AcrIF4 was injected into the buffer flow in increasing concentrations (1.85 nM, 55.6 nM, 167 nM, 500 nM, 1.5 uM), and Csy complex-AcrIF4 binding events were recorded in real time. Experiments were conducted at 37°C, in 20 mM HEPES pH 7.5, 100 mM KCl, 1mM TCEP, 0.005% Tween. Data were fit with a model describing Langmuir binding (i.e. 1:1 binding between free analyte and immobilized ligand). Kinetic rate constants were calculated using Biacore evaluation software (GE).

### Liquid culture phage infections

Liquid culture phage infections and growth curves were performed by infecting 140 µl of *P. aeruginosa* 1:100 diluted overnights cultured in LB supplemented with 10 mM MgSO4 and antibiotics and inducer in a 96 well costar plate with 10 µl of phage diluted in SM buffer. These infections proceeded for 24 hours in a BioTek Synergy microplate reader at 37 °C with continuous shaking. After 24 hours, phage was extracted treating each sample with chloroform followed by centrifugation at 21,000 x g for 2 minutes.

### Lysogen acquisition and induction

Overnight cultures of PA14 were subcultured at 1:100 for ∼3 hours (OD_600nm_ = 0.3) in LB supplemented with 10 mM MgSO4. 1 mL of cells were then infected with 10 µl DMS3m_*acr gp52∷GentR*_ and incubated for 50 minutes at 37 °C, shaking at 100 rpm. The sample was then treated with a 10% volume of 10X gentamicin, spun down at 8,000xg, and resuspended in 200 ul of LB with 50 µg/mL gentamicin. 100 ul of sample was then plated (after further dilution, if required) on gentamicin selection plates and incubated at 37 °C. To analyze the lysogens, the resulting colonies were grown for 16 hours in LB + 10 mM MgSO4 (no selection), the supernatants harvested, and serial dilutions spotted onto a lawn of PA14 ΔCRISPR. Crude genomic DNA for PCR analysis was harvested from the lysogens by boiling 10 µl of culture in 0.02% SDS for 10 minutes.

### Lysogen PCR

PCR amplification of 2 µl of crude genomic DNA harvested from lysogens was used to screen for the presence of *DMS3m-gp52* and *gentR* using MyTaq (Bioline) polymerase with MyTaq GC buffer under standard conditions.

## SUPPLEMENTAL FIGURE LEGENDS

**Figure S1.**
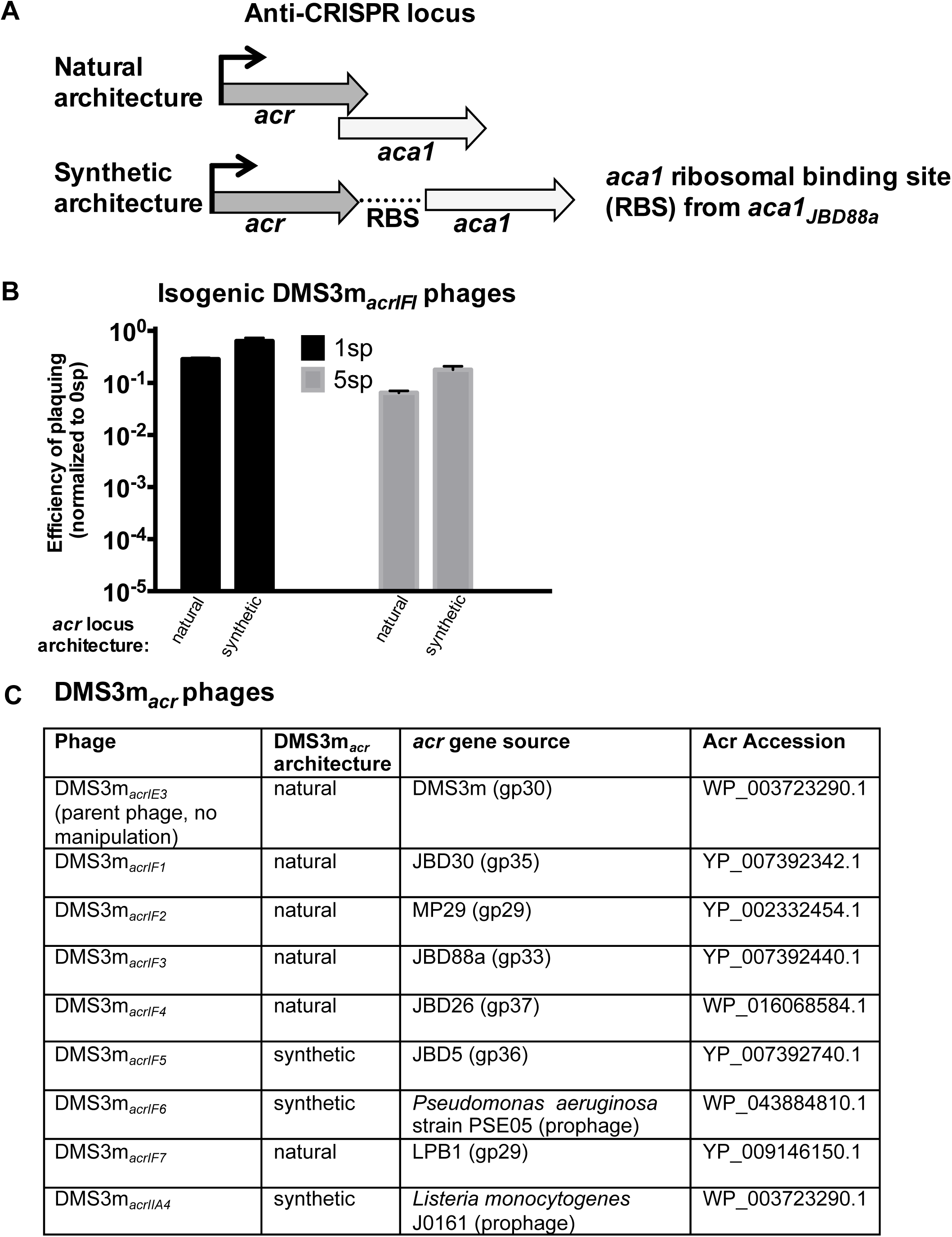
Engineering DMS3m_*acr*_ phages. **(A)** Schematic of two *acr* locus architectures used to generate an isogenic family of DMS3m_*acr*_ phages. **(B)** Efficiency of plaquing (EOP) of DMS3m_*acrIF1*_ with natural or synthetic locus architecture on PA14 with 1 or 5 targeting spacers (1sp and 5 sp). Data are represented as the mean of 3 biological replicates +/-SD. **(C)** Table of DMS3m_*acr*_ phage genotype and locus architecture, as well as the original source of the *acr* gene and the corresponding accession number of the Acr protein.

**Figure S2.**
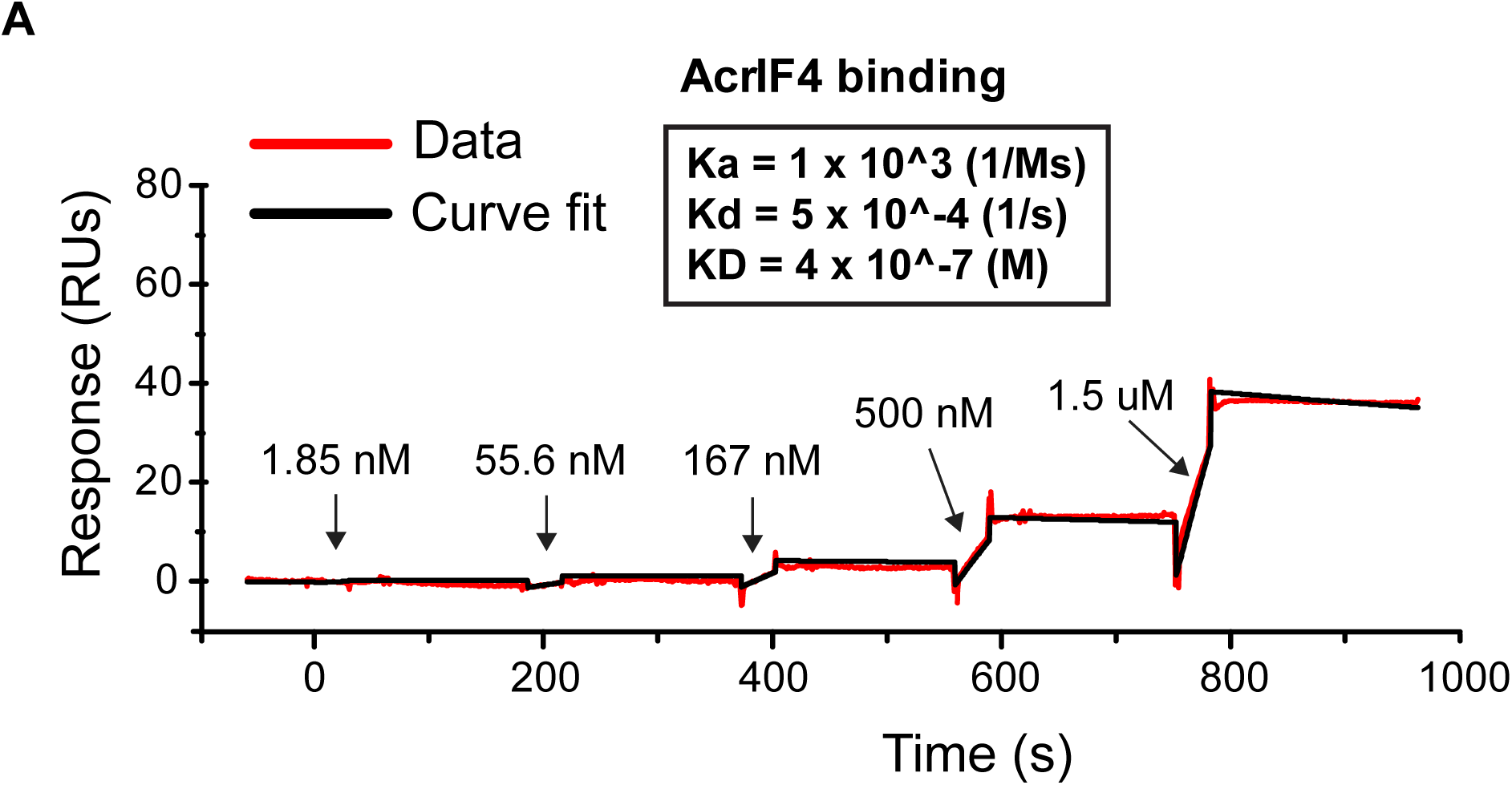
Sensograms of AcrIF4 and AcrIF1 binding the Csy complex. **(A)** Sensogram showing real-time binding of increasing concentrations of free AcrIF4 (1.85 nM, 55.6 nM, 167 nM, 500 nM, 1.5 µM) to immobilized Csy complex. A model describing Langmuir binding (black line) was fit to the data to calculate binding constants (Ka, Kd, and KD; boxed inset). **(B)** Sensogram showing real-time binding of increasing concentrations of free AcrIF1 (1.85 nM, 55.6nM, 167 nM, 500 nM, 1.5 µM) to immobilized Csy complex. A model describing binding at a 2:1 stoichiometry of free analyte:immobilized ligand (black line) was fit to the data to calculate binding constants (Ka1, Ka2, Kd1, Kd2, and KD1, KD2; boxed inset) (Chowdhury and et al. 2017).

**Figure S3.**
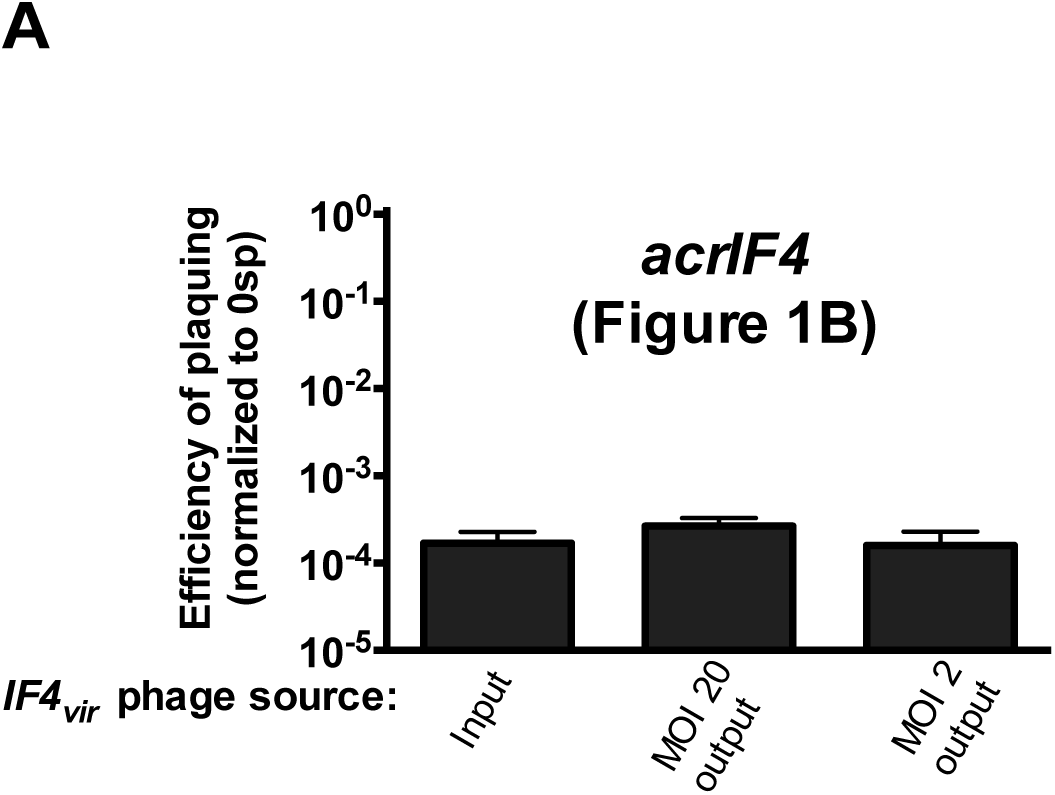
Output phages from Figure 1B are CRISPR sensitive. Efficiency of plaquing (EOP) of the original stock of virulent DMS3macr_*IF4*_ (input) on PA14 5 sp compared to EOP of DMS3m_*acrIF4*_ harvested from high MOI infections (MOI 2, MOI 20 output) from figure 2B. Data are represented as the mean of 3 biological replicates +/-SD.

**Figure S4.**
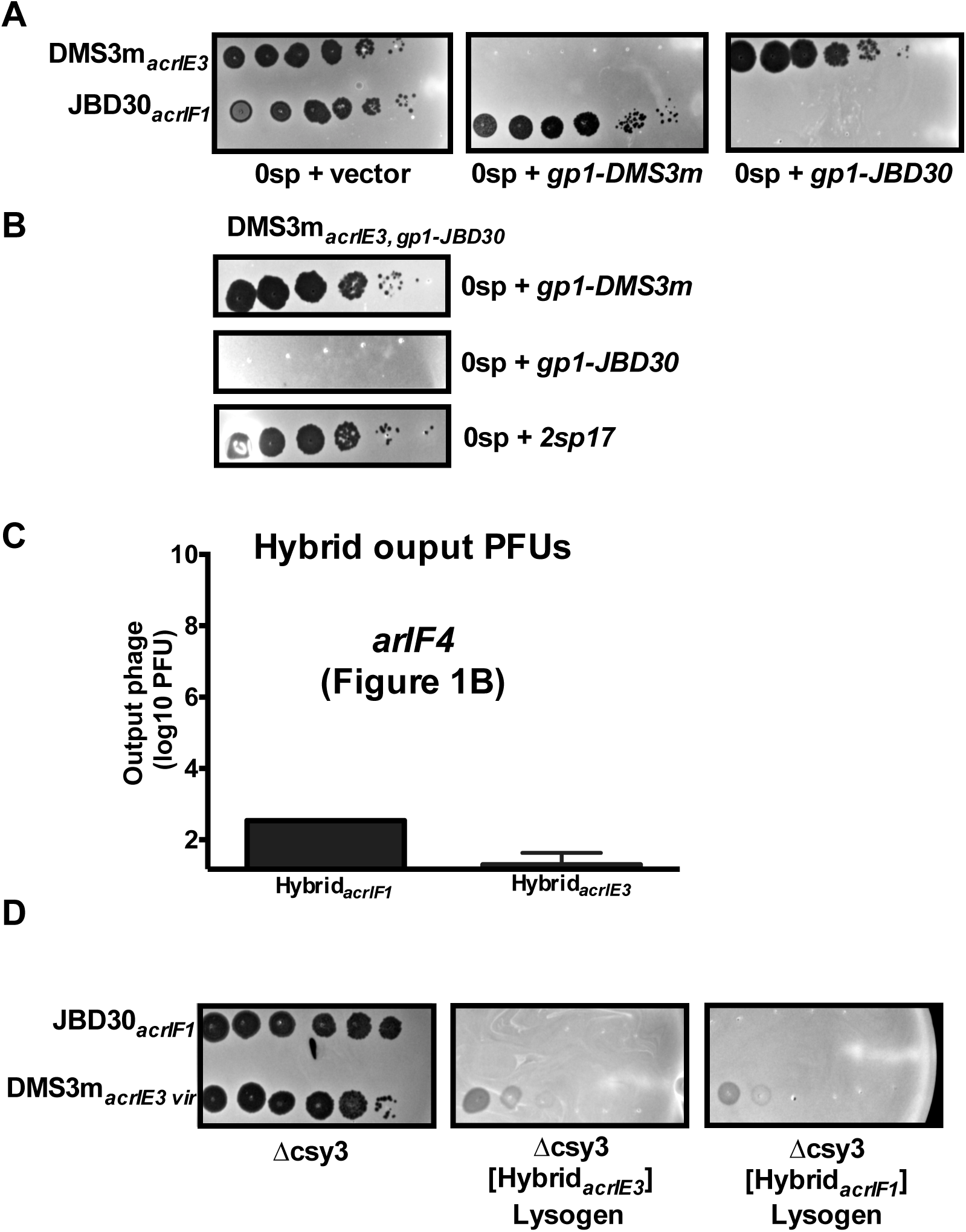
Generating and validating Hybrid_*acr*_ phages. **(A)** 10-fold serial dilutions of DMS3m_*acrIE3*_ and JBD30_*acrIF1*_ spotted on lawns of non-targeting (0sp) *Pseudomonas aeruginosa* PA14 expressing the DMS3m C repressor (*gp1-DMS3m*), the JBD30 C repressor (*gp1-JBD30*) or a vector control. **(B)** 10-fold serial dilutions of hybrid DMS3m_*acrIE3* *gp1-JBD30*_ plated on lawns of non-targeting (0sp) *Pseudomonas aeruginosa* PA14 expressing the DMS3m C repressor (*gp1-DMS3m*), the JBD30 C repressor (*gp1-JBD30*), or a spacer which uniquely targets JBD30 (2sp17). **(C)** Hybrid phage (Hybrid_*acrIF1*_ or Hybrid_*acrIE3*_) harvested from infections of PA14 5sp expressing the JBD30 C repressor from experiments shown in Figure 2G-I. Hybrid PFUs were quantified on the 0sp PA14 strain. Data are represented as the mean of 3 biological replicates +/-SD. **(D)** 10-fold serial dilutions of JBD30_*acrIF1*_ or virulent DMS3m_*acrIE3*_ spotted on lawns of PA14 Δ*csy3* or PA14 Δ*csy3* lysogenized with Hybrid_*acrIE3*_ or Hybrid_*acrIF1*_. Despite being heteroimmune with respect to DMS3, the DMS3 phage is unable to replicate well on this lysogens due to other superinfection exclusion properties of DMS3.

**Figure S5.**
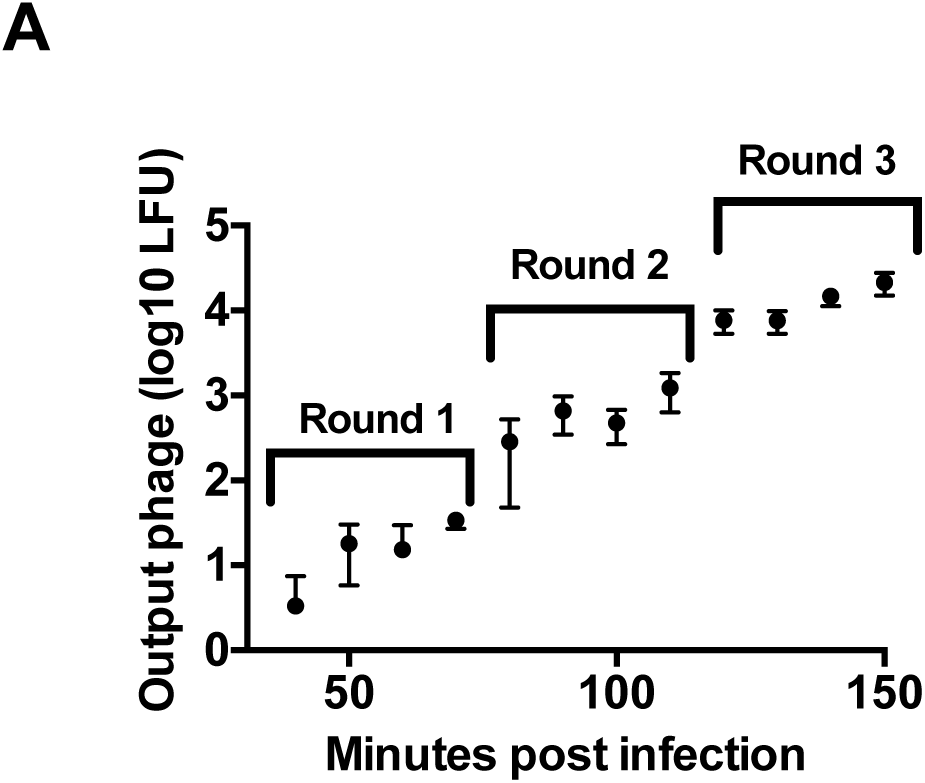
Timecourse of lysogen formation. Time course of acquisition of gentamicin-resistance marked DMS3m_*acrIFI* *gp52∷gentR*_ prophage by a non-targeting (0 sp) strain of PA14, labeled with estimates of each infectious cycle (Rounds 1-3). Data are represented as the mean of 3 biological replicates +/-SD.

**Figure S6.**
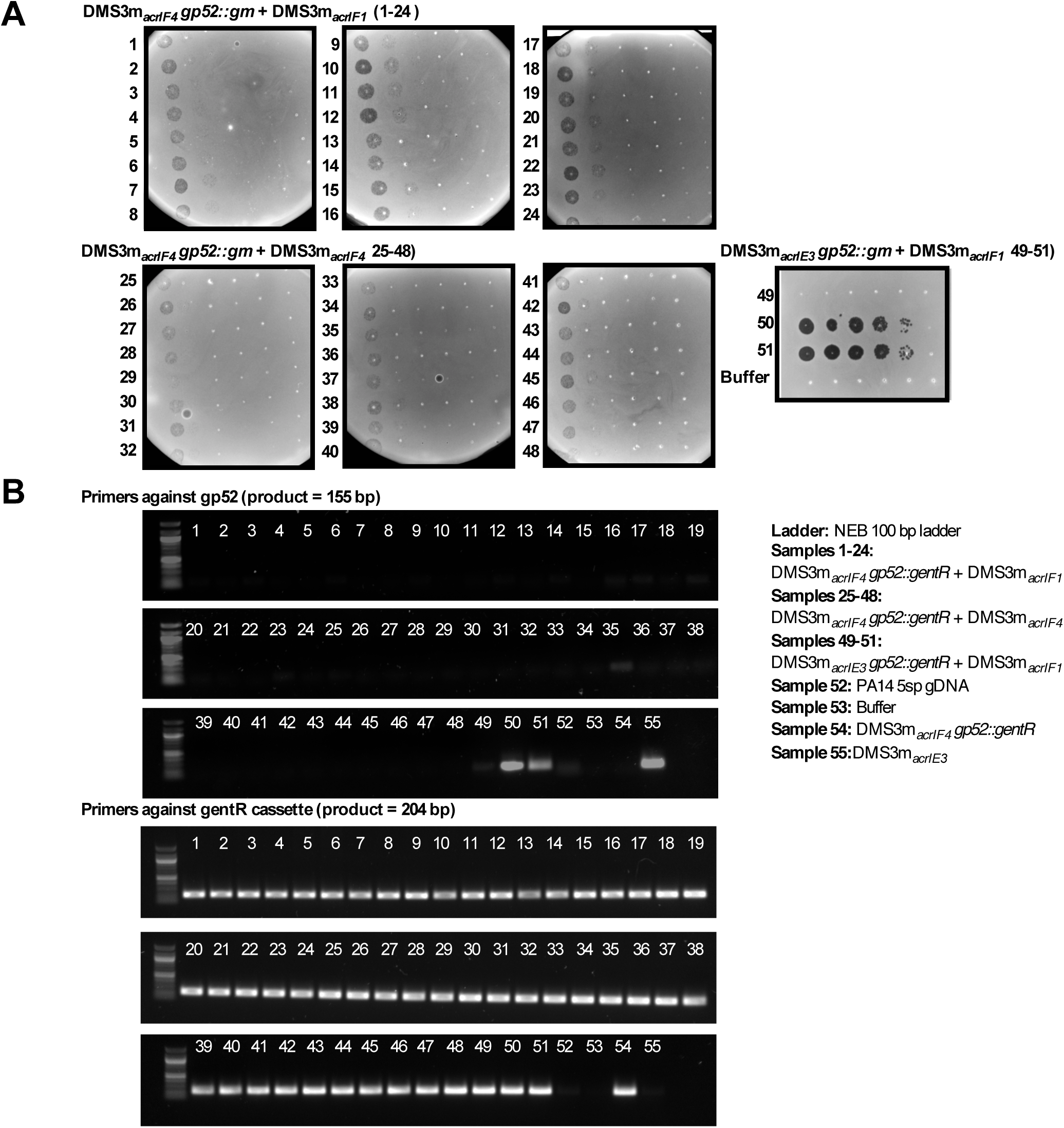
Prophage content of lysogens from Figure 3E-F. **(A)** 10-fold serial dilutions of supernatant harvested from overnight cultures of lysogens from Figure 3E (1-48) and 3F (49-51), spotted on a non-targeting (0 sp) strain of PA14. A faint clearing corresponds to induction of the gentamicin marked phage, while strong plaquing (i.e. 50, 51) reflects the presence of the gentamicin marked phage and the donor phage. **(B)** PCR of genomic DNA harvested from harvested from overnight cultures of lysogens from Figure 3E (1-48) and 3F (49-51) amplified with primers targeting *gp52-DMS3m* (top) or the gentamicin resistance cassette used to replace *gp52* in *DMS3m*_*acr* *gp52∷gentR*_ derivatives (bottom).

**Plasmid table:**

| Plasmid | Backbone | Purpose |
| --- | --- | --- |
| CR2sp17-OE | pHERD 30T | crRNA overexpression |
| CR2sp20-OE | pHERD 30T | crRNA overexpression |
| acrF2-flip | pHERD 30T | generating DMS3m<sub>acrIF2</sub> |
| acrF3-flip | pHERD 30T | generating DMS3m<sub>acrIF3</sub> |
| acrF4-flip | pHERD 30T | generating DMS3m<sub>acrIF4</sub> |
| acrF5-synthetic-flip | pHERD 30T | generating DMS3m<sub>acrIF5</sub> |
| acrF6-synthetic-flip | pHERD 30T | generating DMS3m<sub>acrIF6</sub> |
| acrF7-flip | pHERD 30T | generating DMS3m<sub>acrIF7</sub> |
| acrIIA4-synthetic-flip | pHERD 30T | generating DMS3m<sub>acrIIA4</sub> |
| gp52::gentR-flip | pHERD 20T | generating DMS3m<sub>acr gp52::gentR</sub> |
| gp1-JBD30 | pHERD 20T and 30T | C repressor (JBD30) overexpression |
| gp1-DMS3 | pHERD 20T and 30T | C repressor (DMS3) overexpression |
| sgRNA-DMS3 | pMMBHE | DMS3m targeting with SpyCas9 |
| SpyCas9- pLac | pUC18T-mini-Tn7T-Gm | Insertion of SpyCas9 into the PAO1 tn7 site |

